# Disentangling latent representations of single cell RNA-seq experiments

**DOI:** 10.1101/2020.03.04.972166

**Authors:** Jacob C. Kimmel

## Abstract

Single cell RNA sequencing (scRNA-seq) enables transcriptional profiling at the resolution of individual cells. These experiments measure features at the level of transcripts, but biological processes of interest often involve the complex coordination of many individual transcripts. It can therefore be difficult to extract interpretable insights directly from transcript-level cell profiles. Latent representations which capture biological variation in a smaller number of dimensions are therefore useful in interpreting many experiments. Variational autoencoders (VAEs) have emerged as a tool for scRNA-seq denoising and data harmonization, but the correspondence between latent dimensions in these models and generative factors remains unexplored. Here, we explore training VAEs with modifications to the objective function (i.e. *β*-VAE) to encourage disentanglement and make latent representations of single cell RNA-seq data more interpretable. Using simulated data, we find that VAE latent dimensions correspond more directly to data generative factors when using these modified objective functions. Applied to experimental data of stimulated peripheral blood mononuclear cells, we find better correspondence of latent dimensions to experimental factors and cell identity programs, but impaired performance on cell type clustering.

**Publication Status:** This pre-print represents the final output of a preliminary research direction and will not be updated or published in an archival journal. We are happy to discuss future directions we believe to be promising with any interested researchers.

## 1 Introduction

scRNA-seq experiments can capture many sources of biological variation, including differences between cell identities, responses to perturbations, and developmental programs [1, 2, 3]. The atomic units of an RNA-seq experiment are read counts for individual gene transcripts. However, many processes of interest in biology involve the interaction of many genes in coordinated gene expression programs (GEPs) and may be better represented using a smaller number of dimensions. Many dimensionality reduction methods have been proposed for scRNA-seq data [4, 5, 6, 7], including the recent introduction of variational autoencoder (VAE) based methods [8, 9, 10]. While VAEs have several desirable properties, the latent spaces they learn to encode may be difficult to interpret.

Recent work on VAEs has attempted to encourage “disentangled” latent spaces which may be more interpretable. A disentangled latent space has direct correspondence between dimensions in the latent space and generative factors – parameters of the underlying process that generated the observed data [11]. In the case of single cell RNA-seq data, we may imagine that cell identity, cell cycle state, and the activity of other gene expression programs constitute generative factors. Methods to enforce disentanglement in VAEs largely focus on modifying the objective function [12]. Here, we explore using one of these disentanglement techniques (*β*-VAE) to encourage disentanglement in VAE latent spaces learned for scRNA-seq data. Leveraging simulated data where ground truth values for generative factors are known, we find that these methods improve the correspondence between dimensions of the latent space and generative factors.

### 1.1 Variational Autoencoders

Variational autoencoders (VAEs) learn a generative model of observed data **X** by taking advantage of a lower dimensional latent space **Z**. Briefly, VAEs jointly learn to map observations *x* ∈ **X** to points *z* ∈ **Z** (“encoding”) and to perform the inverse mapping (“decoding”). This two-way mapping is enabled by a flexible encoder *q*(*z|x*) and decoder *p*(*x|z*), often implemented as neural networks. The encoder posterior *q*(*z|x*) is regularized to match a prior distribution on the latent variables, *p*(*z*).

VAEs are trained by jointly optimizing (1) the reconstruction error of observations *x* and (2) divergence of the latent distribution *q*(*z|x*) from the prior *p*(*z*). The VAE objective is traditionally formulated with two components, each addressing one of these desired properties [13].

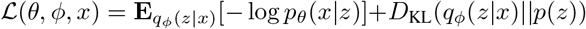

Models of this form have recently been applied to single cell RNA-seq data by multiple groups, yielding effective approaches for gene expression denoising, visualization, and data harmonization [8, 9, 10].

### 1.2 Encouraging disentanglement in the VAE latent space

A standard choice for a Gaussian prior on the latent space 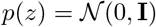 promotes independence among the latent dimensions, since the covariance matrix is diagonal. The *β*-VAE approach [14] leverages this property to enforce independence between dimensions of the latent distribution and encourage disentanglement. This is achieved by weighting the KL-divergence between the prior and latent posterior in the standard VAE objective by a coefficient *β* > 1.

An addition to the *β*-VAE framework encourages an explicit divergence from the prior distribution that increases during the course of training [15]. This is implemented by penalizing the difference between a “channel capacity” constant *C* and the KL-divergence. Together these modifications add two additional parameters to the objective function:

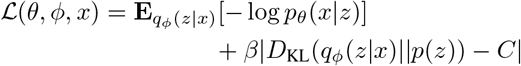

## 2 Experiments

Here, we use the recently introduced Single Cell Variational Inference (scVI) framework[9] as a baseline model, and augment the standard objective function during training by either (1) altering the *β* parameter and/or (2) increasing the channel capacity parameter *C* to encourage disentanglement in the latent space. In this parameter space, *β* = 1 and *C* = 0 represents a baseline scVI model.

For all experiments, we use *n* = 128 units in both the encoder and decoder layers of scVI. We set the number of latent dimensions to 32 and train for 400 epochs using the Adam optimizer with a learning rate of 10^*−*4^ followed by 200 epochs with a learning rate of 10^*−*5^. For experiments where *C >* 0, we linearly increase the channel capacity from *C* = 0.1 to the maximum value over 20, 000 iterations. We fit models for 10 random starts and average evaluation metrics to account for stochasticity in optimization.

### 2.1 Simulated Data

We simulate an scRNA-seq experiment using the Splatter statistical framework [16, 17]. We simulate 10,000 cells with 25,000 genes from 5 cell types. Each cell type is defined by expression of a “cell identity” GEP. We also simulate an “activity” GEP which is utilized within 3 of the 5 cell types (Fig. 1A).

**Figure 1:**
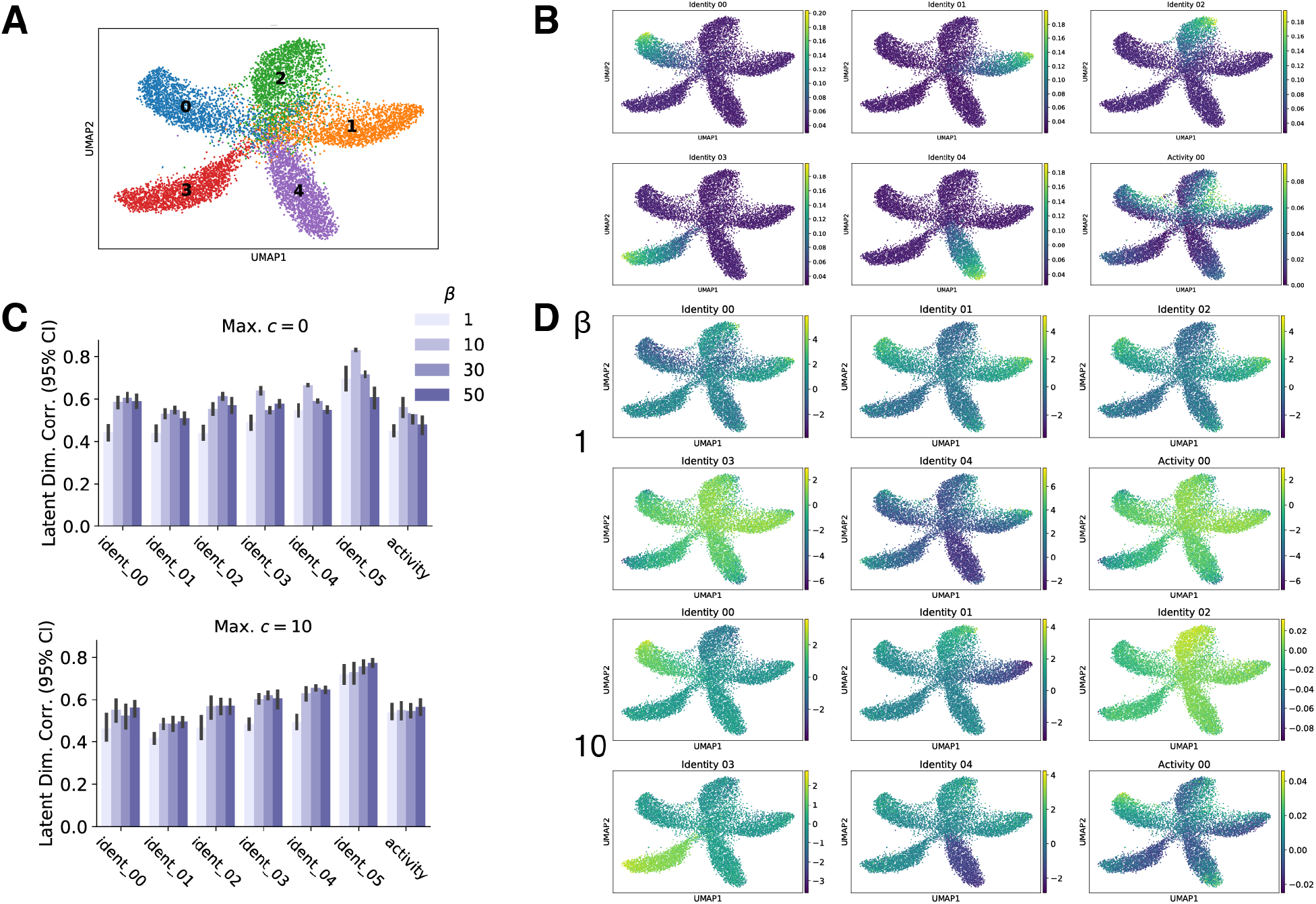
Increasing *β* improves correspondence between simulated GEPs and VAE latent dimensions. **(A)** UMAP projection of simulated single cell data. Ground truth cell type identities for each cell are overlaid in color. **(B)** Ground truth rank-based GEP score of the 5 cell type identity programs and the “activity” program in each cell. **(C)** Maximum absolute Spearman correlation of GEP scores with dimensions of the latent space as a function of *β*. **(D)** Values for the latent dimension with the highest correlation with each GEP are presented on a UMAP projection.

This dataset provides a ground truth (1) cell identity, and (2) gene sets associated with each GEP. Formally, we define a GEP as a group of genes that are co-differently expressed. Each gene in a GEP is scaled by a differential expression coefficient *D* where 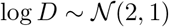 in cells where that GEP is active. To quantify GEP utilization in each cell, we compute a rank-based score in the same manner as AUCell [18] that we refer to as a “GEP score”. We consider the scores of these ground truth GEPs to be generative factors in the data which we wish to capture in dimensions of the latent space (Fig. 1B).

### 2.2 Recovering generative factors in simulated data

To determine if a modification to the VAE objective improves interpretability of the latent space, we require a quantitative metric for interpretability. Quantitative metrics for disentanglement are still an active area of research and no well-defined standard exists [12]. Here, we focus on a simplistic metric to evaluate the correspondence between latent dimensions and ground truth GEPs. For each ground truth GEP, we compute Spearman correlations *ρ* of the GEP score with each dimension of the latent space. We consider the maximum absolute correlation to reflect the best correspondence between a latent dimension and a GEP. This metric does not reflect overall disentanglement in the latent space, which inherently must consider the uniqueness of correspondence between latent dimensions and generative factors.

We fit scVI models with varying values for *β* and *C*. We find that increasing *β* > 1 while holding *C* = 0 significantly increases the correlation between ground truth GEPs and latent dimensions at some values (*β* = 10, *t*-test, *q* < 0.05, Benjamini-Hochberg). However at higher values (*β* = 50), decreased performance is observed for some ground truth GEPs (Fig. 2A). Increasing the channel capacity to *C* = 10 ameliorates the detrimental effect from larger values of *β*, although the maximum correlation between some GEPs and a latent dimension is decreased (Fig. 1C).

**Figure 2:**
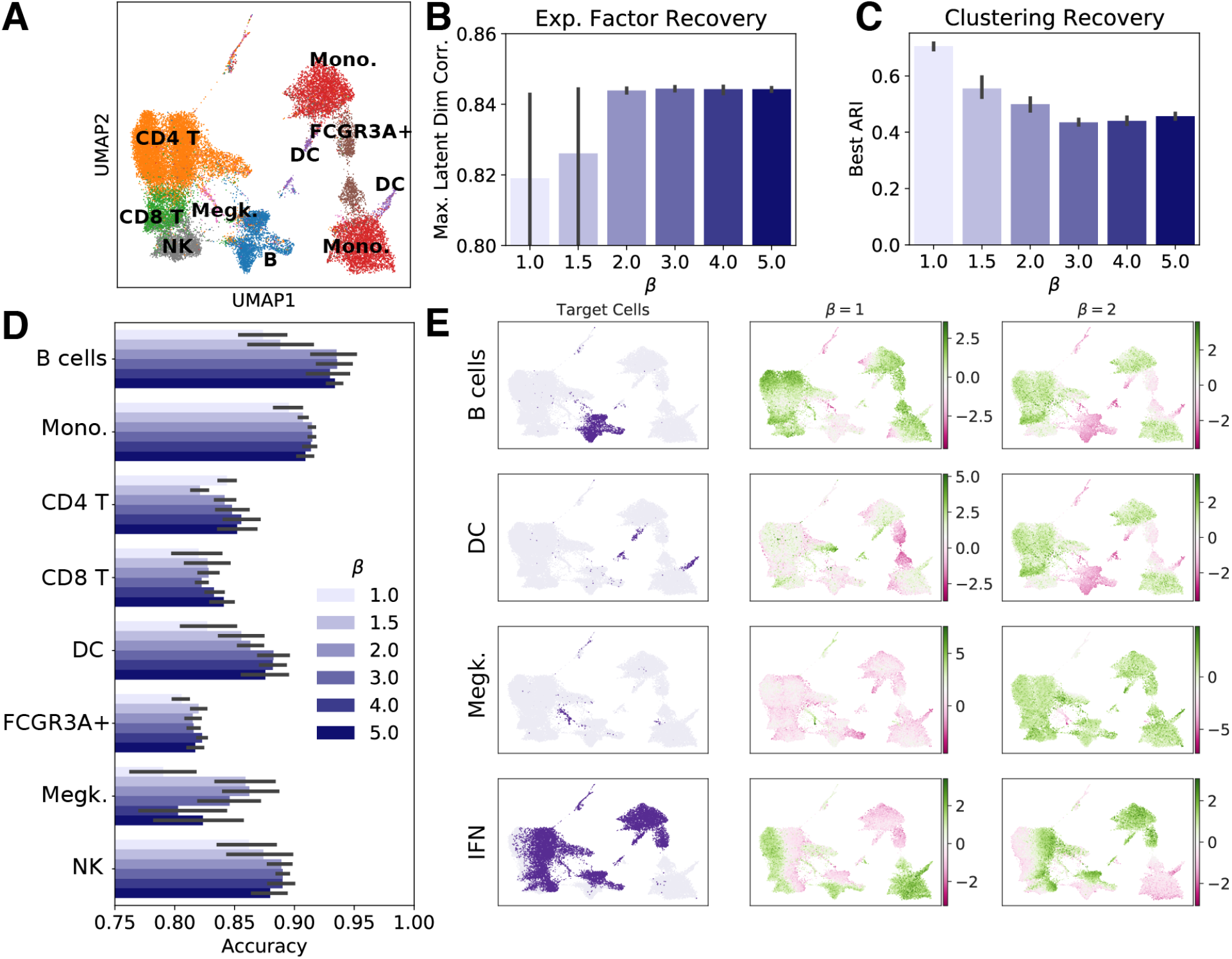
Increasing *β* modestly improves experimental factor recovery but impairs cell type clustering. **(A)** UMAP projection of PBMC data with cell type labels. **(B)** Correlation of latent dimensions with IFN*β* treatment status. **(C)** Adjusted Rand Index for cell type recovering by community detection in latent spaces. **(D)** Max logistic regression cell type classification accuracy (mean 5-fold CV) based on a single latent dimension. **(E)** Values of the latent dimension with the best correspondence to each cell identity program or IFN*β* condition (rows) are visualized in UMAP projections. Cells that should be distinguished by each latent dimension are highlighted on the left, and the best dimension from VAEs with *β* = 1 (center) or *β* = 2 (right) are shown.

To determine if highly correlated latent dimensions correspond to specific ground truth GEPs, we visualize the values of the latent dimension most correlated with each ground truth GEP in a UMAP projection. Using the baseline *β* = 1, we find that latent dimensions are not specific for a ground truth GEP. Using *β* = 10 appears to improve the correspondence of latent dimensions to ground truth GEPs. We note that with *β* = 10, the most correlated latent dimensions for each cell identity GEP appear to specifically mark that cell identity (Fig. 1D). For the latent space learned with *β* = 10, note that the dimensions for identities 1 and 5 are inverse mappings and that the dimension for identity 2 shows a less dramatic correspondence than other GEP:dimension pairs.

### 2.3 Recovering experimental perturbations and cell identity in PBMCs

To determine if modified VAE objectives can capture cell identity programs and experimental factors, we trained scVI models on experimental data from peripheral blood mononuclear cells (PBMCs) [19]. The data contain 8 unique cell types, each of which is observed before and after stimulation with IFN*β* (Fig. 2A). We treat the IFN*β* experimental condition as a generative factor we wish to recover. As before, we evaluate the maximum Spearman correlation between latent dimensions and IFN*β* condition for a range of *β*. We find that increasing *β* leads to modest improvements in this correlation, but even *β* = 1 models learn a dimension with strong correlation (Fig. 2B). Visualizing the latent dimensions that correspond best to IFN*β* condition confirms that even the *β* = 1 models learn a dimension that segregates the experimental conditions (Fig. 2E).

Latent spaces are also used for unsupervised cell type identification by Louvain community detection [20]. We evaluate this method in each latent space using the Adjusted Rand Index (ARI) between the Louvain partition and ground truth cell types. We use a range of resolutions and take the maximum ARI to mimic human adjustment based on visualization. We find that unsupervised clustering efficacy decreases as *β* increases, suggesting that stronger regularization may be undesirable for some downstream tasks (Fig. 2C).

To determine if we recover cell identity GEPs in each latent space, we fit logistic regression models to classify each cell type based on each individual latent dimension. We fit each regression model to distinguish a single target cell type (i.e. B cells) from all other cell types. We perform class balancing before fitting and report accuracy as the mean of 5-fold cross-validation. Here, we assume that a latent dimension representing a cell identity program will allow for better classification of the corresponding cell type. We consider the dimension with the maximum classification accuracy to have the best cell identity program correspondence.

We find that increasing *β* > 1 improves the correspondence of latent dimensions and cell identity programs by this metric for most cell types (Fig. 2D, E). However, counterexamples also exist – we find that correspondence between latent dimensions and the CD4 T cell identity program decreases for some values of *β* > 1. Taken together, these results suggest that modifying the VAE objective can improve correspondence between latent dimensions and some generative factors, but may also decrease performance on some downstream tasks like cell type clustering.

## 3 Conclusions

We find that a modified VAE objective (*β*-VAE) designed to encourage disentanglement improves the correspondence between VAE latent dimensions and ground truth gene expression programs in simulated data. Applied to experimental data, we observe modest improvements in the correspondence of latent dimensions with experimental conditions and cell identity programs. However, we also find that the performance of cell type clustering decreases in the same conditions.

These results suggest that fitting VAE models to scRNAseq data with the *β*-VAE objective may improve the interpretability of latent spaces, but that these modifications may decrease performance on some downstream tasks. We note that the simplistic metrics of correspondence we employ here do not measure disentanglement directly. We believe these results motivate further research on the application of disentanglement methods to single cell RNA-seq models. Multiple groups have proposed alternative formulations of the VAE objective that may ameliorate the detrimental effects of disentanglement methods we observe here. Likewise, additional quantitative metrics of disentanglement have been proposed that may more accurately identify alignment of latent variables with generative factors [21, 22, 23]. Application of these techniques may prove fruitful and remains an exciting direction for future work.

